# Analysis of *Treponema pallidum* strains from China using improved methods for whole-genome sequencing from primary syphilis chancres

**DOI:** 10.1101/2020.04.15.043992

**Authors:** Wentao Chen, David Smajs, Yongfei Hu, Wujian Ke, Petra Pospíšilová, Kelly L. Hawley, Melissa J. Caimano, Justin D. Radolf, Arlene Sena, Joseph D. Tucker, Bin Yang, Jonathan J. Juliano, Heping Zheng, Jonathan B. Parr

**Author notes:** co-senior authors (Heping Zheng, Jonathan B. Parr). **Corresponding authors:** Jonathan B. Parr, MD, MPH, University of North Carolina, 130 Mason Farm Rd, Chapel Hill, NC, USA. Heping Zheng, PhD, Dermatology Hospital, Southern Medical University, N0.2 Lujing Rd, Guangzhou, P. R. China.

## Abstract

Whole-genome sequencing (WGS) of *Treponema pallidum* subsp. *pallidum* (TPA) has been constrained by the lack of *in vitro* cultivation methods for isolating spirochetes from patient samples. We built upon recently developed enrichment methods to sequence TPA directly from primary syphilis chancre swabs collected in Guangzhou, China. By combining parallel, pooled whole-genome amplification (ppWGA) with hybrid selection, we generated high quality genomes from four of eight chancre-swab samples and two of two rabbit-passaged isolates, all subjected to challenging storage conditions. This approach enabled the first WGS of Chinese samples without rabbit passage and provided insights into TPA genetic diversity in China.

## INTRODUCTION

Efforts to address the resurgence of the sexually transmitted disease syphilis have been hampered by the limited understanding of the genetic diversity of its causative pathogen, the spirochete *Treponema pallidum* subsp. *pallidum* (TPA) [1]. A vaccine to prevent TPA infection and transmission is sorely needed. Design of an effective syphilis vaccine requires an understanding of the diversity of TPA strains circulating worldwide, particularly in countries such as China, where the incidence of disease has steadily increased by more than 30-fold since the mid-1990s but from which only nine TPA genomes have been reported to date [2–5].

Genomic analyses of TPA have lagged behind those of other pathogens, largely due to the lack of *in vitro* methods for isolating these spirochetes from patient samples [6]. Since publication of the first complete TPA genome in 1998, only a small number of TPA genomes have been published, and the majority of sequenced isolates required passage in rabbit testicles prior to sequencing [7]. Several strategies have been used to enrich TPA from clinical specimens. Selective enrichment of TPA DNA through hybridization with RNA oligonucleotides for pull-down of DNA fragments with homology to known target sequences has now been employed in several recent TPA genomic analyses [8–10]. Two newer techniques have also been described, one that utilizes anti-treponemal antibody binding to enrich TPA cells isolated from clinical specimens and another that employs methyl-directed enrichment using the restriction nuclease *DpnI* [11, 12]. While these methods represent advancements for the field, improved and complementary approaches for sequencing TPA in patient samples are needed, especially for samples with low TPA burdens or DNA loss during sample processing.

In the present study, we build upon recent advances in TPA enrichment to expand the range of samples from which whole-genome sequences can be obtained. We successfully sequenced TPA DNA extracted from both rabbit-passaged isolates and chancre swabs from Guangzhou, China. Using a combination of ppWGA and hybrid selection, we achieved more than 80% genomic coverage in a majority of samples tested, despite challenging sample processing and storage conditions. Phylogenomic analyses revealed diverse TPA strains, including the first Nichols-like genome published from China to date.

## METHODS

### Study population and sample processing

We collected samples from chancre exudates in Guangzhou, China, from 2017 to 2018. In brief, genital ulcers of patients diagnosed with primary syphilis were swabbed and processed as described in the **Supplementary Methods.** For the present study, we selected 10 samples from male patients ages 20 to 68 for sequencing (**Supplementary Table 1**). These included eight DNA samples extracted directly from chancre swabs and two samples extracted from rabbit testes after intratesticular passage. This study was approved by the ethics committee of the Dermatology Hospital of Southern Medical University (GDDHLS-20170614). The University of North Carolina at Chapel Hill (UNC) institutional review board determined that the analysis of de-identified TPA DNA samples did not constitute human subjects research (UNC study #18-1949). All patients provided written informed consent and were offered treatment as part of routine care.

TPA genome copy numbers and total DNA concentrations were quantified using real-time quantitative PCR (qPCR) and a Qubit 4.0 fluorimeter with dsDNA HS reagents (Thermo Fisher Scientific, Waltham, Massachusetts, USA), respectively, prior to freeze-drying and overnight shipment to UNC at ambient temperature (see **Supplementary Methods**). Freeze-drying was performed in order to comply with regulatory requirements and to facilitate long-distance shipment. Dehydrated samples were stored at −80 °C until resuspension in 25 μl or 100 μl of Buffer EB (QIAGEN, Venlo, Netherlands).

### ppWGA, hybrid selection, and library preparation

Total DNA concentrations of resuspended samples were quantified using a Qubit fluorimeter as described above. For those samples with concentrations below the fluorimeter’s limit of detection (0.2 ng/ul), whole-genome amplification (WGA) using the Illustra GenomiPhi V2 Amplification kit (GE Healthcare, Chicago, Illinois, USA) was employed to increase the library input. Three μL of DNA template was used for each reaction; reactions were otherwise performed according to the manufacturer’s instructions with the exception of reaction time, which was increased to 6 hours due to the low DNA concentration. Five separate WGA reactions were performed in parallel for each individual sample. The amplification products for each sample were pooled to achieve equal total DNA input per WGA reaction and purified using KAPA Pure Beads (Kapa Biosystems, Wilmington, Massachusetts, USA).

TPA DNA was enriched using the SureSelect XT HS target enrichment system (Agilent Technologies, Santa Clara, CA, USA), which utilizes RNA oligonucleotide probes to hybridize DNA of interest. We custom-designed probes using all TPA genomes that were publicly available at the time of construction, with increased tiling density of probes across specific regions of interest, including phylogenetically informative loci and those that encode known or putative outer membrane proteins (**Supplementary Tables 2-3**) [13]. Probe design, library construction, and hybridization were performed as described in the **Supplementary Methods**. Sequencing was performed at the UNC High-Throughput Sequencing Facility using the MiSeq platform (Illumina, San Diego, CA) with 150 bp paired-end reads.

### Genomic alignment, variant calling, and phylogenetic analysis

For phylogenetic analysis, we included 68 publicly available (18 complete and 50 draft), geographically diverse TPA genomes from three continents, in addition to the genomes generated in the present study. Sequencing alignment, variant calling, and phylogenetic analysis were performed as depicted in **Figure 1**. Putative sites of recombination and indels were removed prior to phylogenetic analysis. We assessed the validity of our variant calls and phylogenetic analysis by Sanger sequencing five targets, including three loci used in a multilocus strain typing (MLST) system recently proposed by Grillova *et al.* (*tp0136*, *tp0548* and *tp0705*) and each of TPA’s two 23S rRNA operons, mutations in which have been associated with azithromycin resistance (see **Supplementary Methods**) [14]. Sequences were uploaded to the Sequence Read Archive and GenBank (accession numbers pending).

**Figure 1.**
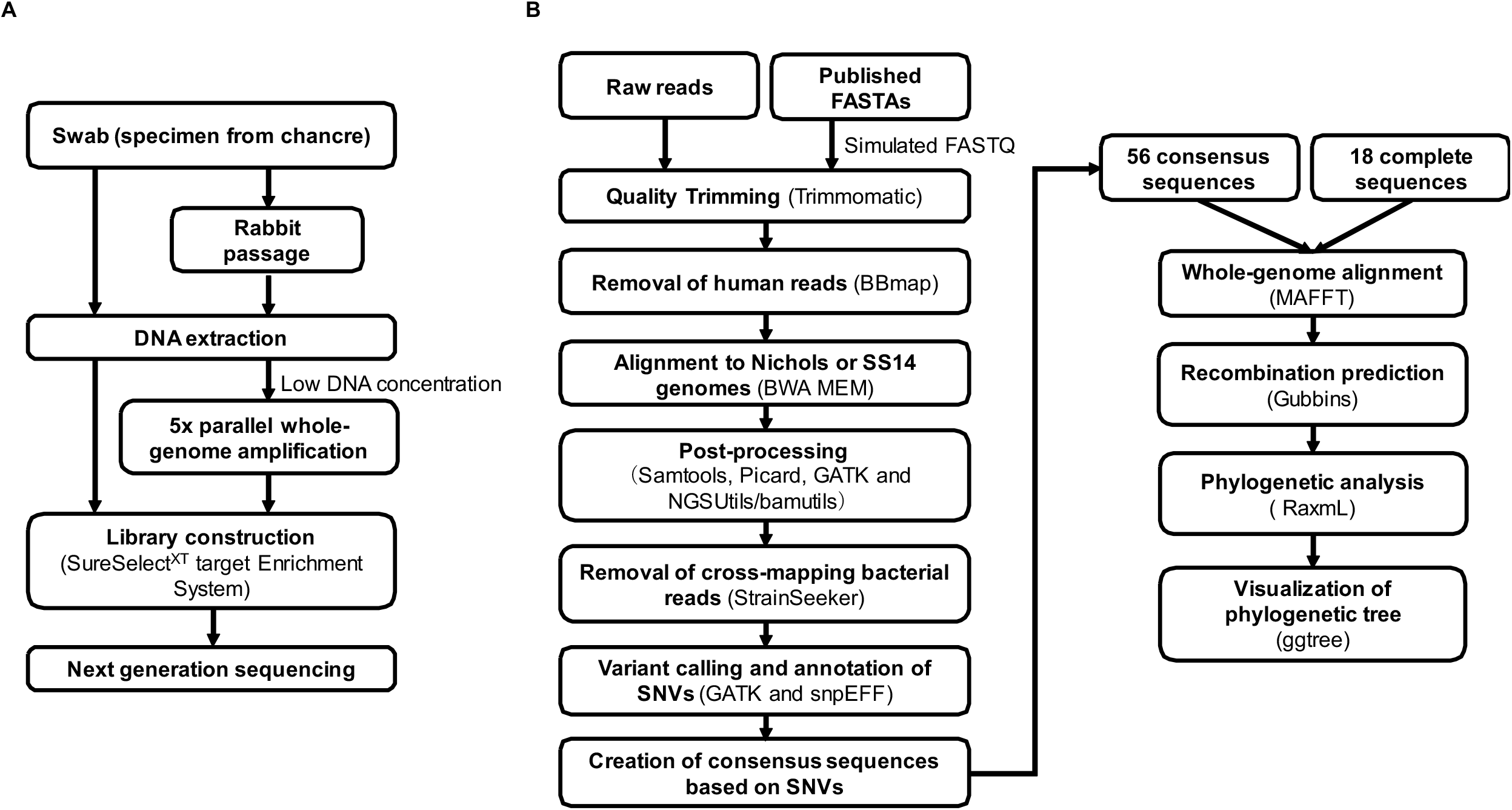
(A) Sample enrichment, sequencing library preparation, and (B) bioinformatic processing. DNA was enriched for TPA prior to sequencing using rabbit passage; five individual, parallel WGA reactions that were then pooled (ppWGA); and/or hybrid selection with RNA baits. (B) Sequences were aligned to the Nichols or SS14 reference genomes prior to variant calling and phylogenomic analysis.

## RESULTS

### Whole-genome sequencing outcomes

After freeze-drying and rehydration, only three samples had sufficient DNA concentrations for quantification by Qubit. Total DNA concentrations for these samples ranged from 1.3 to 10.6 ng/uL, representing a 47 to 58% decrease in total DNA compared to the original samples (**Supplementary Table 4**). The remaining seven rehydrated DNA samples had concentrations beneath the fluorometer’s limit of detection and were subjected to ppWGA before hybridization (**Figure 1**). Despite evidence of DNA loss during the process of freeze-drying, shipment, and rehydration, we successfully sequenced TPA genomes from six of 10 samples (60%), including two of two rabbit-passaged isolates and four of eight chancre-swab samples that had not undergone rabbit passage. Among the six samples with sufficient coverage for phylogenomic analysis, 83.5 to 99.1% of the genomes were covered with ≥3 reads.

Sequencing coverage roughly correlated with the original TPA concentrations before freeze-drying (**Supplementary Table 4**). Samples that originally contained 3,320 to 3,160,000 copies of *polA* per μL before freeze-drying achieved ≥3x depth in >80% of their genomes. Sequences from samples subjected to ppWGA before hybridization demonstrated evidence of significant “jackpotting” events (**Figure 2A**) –very high coverage in specific regions – some of which were unique to a given sample, while others were shared across samples. Although these jackpotting events decreased the cost efficiency of the sequencing runs, the use of ppWGA reactions nonetheless enabled salvage of low-concentration, rehydrated samples. In the two samples that were subjected to both approaches (SMUTp_08 and SMUTp_09), ppWGA achieved more even coverage than conventional WGA performed in singleton (**Supplementary Figure 1**). Together, ppWGA and hybrid selection allowed us to generate the first Chinese TPA genomes without rabbit passage.

**Figure 2.**
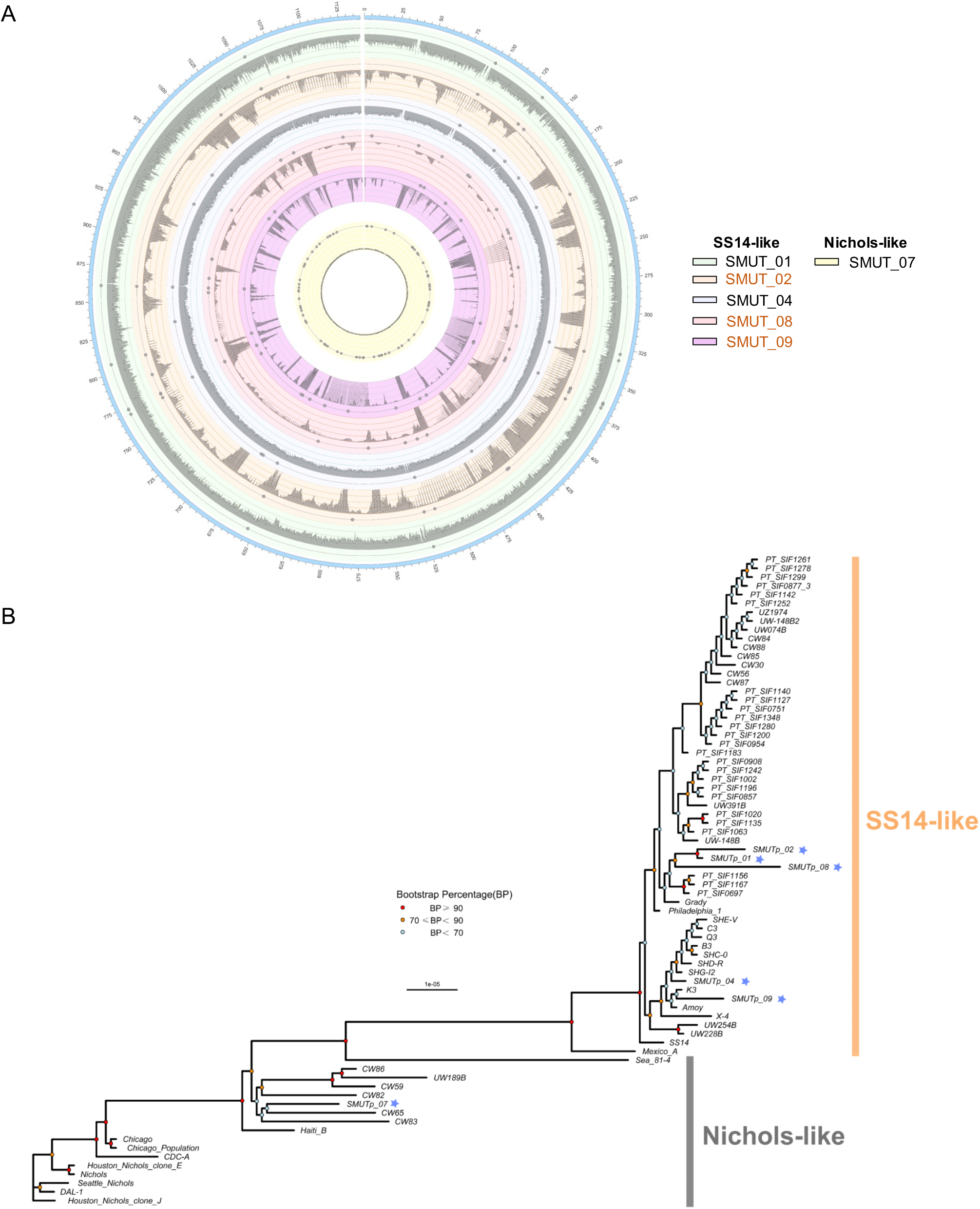
**(A) Genomic coverage and SNVs of new Chinese TPA isolates**, using coordinates from the SS14 or Nichols reference genome. From the outer ring to center, the circles represent six Chinese isolates: SMUTp_01, SMUTp_02, SMUTp_04, SMUTp_08, SMUTp_09, and SMUTp_07, respectively. The grey histogram depicts the depth of sequencing coverage and highlights jackpotting events in samples subjected to ppWGA (labelled in orange, SMUTp_02, SMUTp_08, and SMUTp_09). SNVs are represented by small circles beneath the coverage histogram, filtered for ≥3x depth of coverage. **(B) Phylogenomic diversity of new and published TPA strains**. Maximum likelihood phylogenies were generated from a whole genome alignment of 56 draft and 18 complete TPA genomes, excluding recombinant loci. New Chinese genomes generated in the present study are starred (*).

### Phylogenetic diversity of Chinese TPA isolates

We observed diverse TPA strains within our modest sample set (**Figure 2B**). After exclusion of 66 putative sites of recombination from whole-genome phylogenetic analysis, our sample set included 94 SNVs expected to induce coding changes (**Supplementary Tables 5-9)**. We did not observe obvious clustering of rabbit-passaged TPA isolates, a finding that might be observed if passage in rabbits exerts selective pressure favoring specific variants, versus directly sequenced TPA isolates.

The genomes produced in the present study fell into three distinct clusters (**Figure 2B**) and could be distinguished from the previously published Chinese Amoy genome (**Supplementary Table 10**). These clusters included one closely related to previously sequenced Chinese SS14-like isolates, one distinct SS14-like cluster, and one Nichols-like genome (SMUTp_07) [2–5, 9, 10]. MLST results confirmed the strain relationships observed during phylogenomic analysis (**Supplementary Table 11**). While past surveys using traditional molecular strain typing methods indicated low-level prevalence of Nichols-like strains in China [9, 15], this is the first Nichols-like whole-genome sequence reported from China to our knowledge. Furthermore, this finding has implications for vaccine design, due to the possibility of exchange of genetic material between co-circulating Nichols- and SS14-like strains in a population.

Initial whole-genome alignments of sample SMUTp_02 suggested the presence of a strain with one wild-type (macrolide-sensitive) 23S rRNA operon. However, manual inspection of these reads revealed poor mapping and sequences with 100% homology to *Pseudomonas* species. Sanger sequencing revealed that all samples harbored mutations associated with macrolide resistance in both 23S rRNA operons (**Supplementary Table 12**).

## DISCUSSION

The lack of an *in vitro* cultivation system for TPA and the need for rabbit passage have limited syphilis research. Recently developed sample enrichment and sequencing techniques offer new opportunities to study TPA transmission and evolution. However, they often fail when applied to clinical samples isolated directly from patients due to low TPA burdens. While the number of publicly available TPA genomes has increased rapidly in the past several years, most published genomes required TPA enrichment by rabbit passage prior to WGS.

We built upon recent advances in TPA genomics by piloting a novel WGA approach (ppWGA) for enrichment of challenging clinical samples with low DNA concentrations prior to hybrid selection. Enrichment success using hybrid selection alone has previously been achieved with clinical samples with more than 1 × 10^4^ TPA copies/μL [8]. However, this approach is expected to fail for many clinical samples collected directly from patients, which often harbor only 10^1^ - 10^4^ TPA copies/μL. The success of our approach in samples with concentrations nearing 10^3^ TPA copies/μL measured prior to demanding sample processing and shipment confirms the utility of ppWGA as an adjunct to other TPA enrichment methods. These methods have potential for use in other fields and with diverse sample types, especially when freeze-drying is required for regulatory compliance or samples have low target concentrations.

WGA is typically employed in singleton for non-specific pre-amplification prior to selective enrichment and/or sequencing library preparation. During pilot testing in the present study, we observed distinct, isolated regions of deep sequencing coverage (“jackpotting”) in samples subjected to a single WGA. These findings may be due to early priming by random hexamers and amplification during the WGA process. To overcome the apparent stochastic nature by which genomic locations undergo jackpotting during a WGA reaction, we used WGA in parallel and pooled products before proceeding with hybrid selection and library preparation.

Our study is limited by its small sample size and the challenges of low-concentration and degraded samples. While ppWGA enabled successful enrichment of a portion of these samples, the presence of *Pseudomonas* in one of the downstream 23S rRNA alignments confirms the need for careful attention to regions that are highly conserved across bacteria during analysis. We overcame the challenge of incorrect mapping of short reads by incorporating rigorous quality filters and removing reads that cross-mapped to other bacteria during bioinformatic analysis. These measures are especially important when non-specific amplification like ppWGA is employed during the sample enrichment process.

We employed ppWGA and hybrid selection to gain new insights into the genetic diversity of TPA strains currently circulating in China, where there is a syphilis epidemic and only a small number of rabbit-passaged isolates have been published [2–5]. Additional studies using novel enrichment methods and in locations where little is known about TPA genomic diversity, including sites outside of the United States and Europe, are needed to inform syphilis vaccine design.

## NOTES

### Acknowledgments

The authors would like to thank Madeline Denton for assistance with TPA WGA experiments, Dr. Jiajian Zhou for assistance with NGS data analysis, and Prof. Tianci Yang for providing plasmid DNA for *polA* qPCR standards.

### Financial support

This study was supported by the Medical Scientific Research Foundation of Guangdong Province, China (A2018264, B2019022 to WTC), Science and Technology Planning Project of Guangdong Province, China (2017A020212008 to HPZ). Additional support was provided by the National Institute for Allergy and Infectious Diseases (U19AI144177 to DS, KLH, MJC, JDR, AS, JDT, BY, JJJ, HZ, and JBP, K24AI143471 to JDT, K24AI134990 to JJJ, R01AI26756 to JDR), a Yang Biomedical Scholars Award to JJJ, the Clara Guthrie Patterson Trust, Bank of America, N.A., Trustee (KLH), the Connecticut Children’s Medical Center (JDR, MJC and KLH), the Grant Agency of the Czech Republic (GA17-25455S; gacr.cz to DS) and an American Society for Tropical Medicine and Hygiene-Burroughs Wellcome Foundation award to JBP.

### Potential conflicts of interest

All authors report no conflicts.

## SUPPLEMENTARY TABLES

All Supplementary Tables are provided in a single, attached file for convenience and ease of viewing.

**Supplementary Table 1. Clinical information for Chinese samples included in this study.** See attached file.

**Supplementary Table 2. Genomes used during the design of RNA oligonucleotide “baits” for hybrid selection.** The SS14 and Nichols genomes were covered with 1x tiling density (one bait per locus) with the exception of loci provided in Supplementary Table 3. See attached file.

**Supplementary Table 3. Genomic regions of increased bait tiling density (5x).** Phylogenetically informative loci and those encoding known or putative outer membrane proteins were covered with at least 5x tiling density (five baits per locus). See attached file.

**Supplementary Table 4. Sample details and sequencing results.** See attached file.

**Supplementary Table 5. Putative regions of recombination identified by Gubbins and excluded from phylogenetic analysis.** See attached file.

**Supplementary Table 6. Called variants among the five Chinese SS14-like TPA isolates described in the present study,** annotated using *SnpEff*, prior to removal of putative sites of recombination and paralogous genes. See attached file.

**Supplementary Table 7. Called variants for the single Chinese Nichols-like TPA isolate described in the present study,** annotated using *SnpEff*, prior to removal of putative sites of recombination and paralogous genes. See attached file.

**Supplementary Table 8. Called variants expected to induce coding changes (amino acid changes) among the six Chinese TPA isolates described in the present study.** See attached file.

**Supplementary Table 9. Mutations in penicillin-associated genes.** See attached file.

**Supplementary Table 10. Comparison of SNVs between the new Chinese SS14-like strains (present study) and the Amoy strain.** See attached file.

**Supplementary Table 11. MLST allelic profiles of Chinese TPA strains.** See attached file.

**Supplementary Table 12. Genetic markers of macrolide (azithromycin) resistance.** See attached file.

**Supplementary table 13. Sequences used during phylogenetic tree constuction.** See attached file.

## Notes

### Competing Interest Statement

The authors have declared no competing interest.

