## Supplementary Materials for "Analysis of *Treponema pallidum* strains from China using improved methods for whole-genome sequencing from primary syphilis chancres"

**[#] Corresponding authors:**

### SUPPLEMENTARY METHODS

*Sample processing*

Lesions were first cleansed using sterile saline, then pressured at the base to increase the exudate. Exudate was collected with a sterile coverslip, which was then inverted and placed on a clean glass slide for darkfield microscopy. Next, remaining exudate was swabbed using a sterile polyester swab (KANGJIAN Medical Apparatus Co. Ltd., China) and immediately placed in storage buffer (85% RPMI-1640 medium + 15% glycerol). Darkfield microscopy was then performed by searching for spirochetes systematically with the 40X objective, and examining identified spirochetes with the 100X objective.

Two-hundred μL of buffer was aliquoted for DNA extraction using QIAamp DNA Mini kits (QIAGEN) according to the manufacturer's instructions. Samples underwent real-time PCR targeting the TPA DNA polymerase I (*polA*) gene, performed as previously described [1]. A subset of PCR-positive samples was passaged in rabbits by intratesticular inoculation of seronegative New Zealand white rabbits following a previously described protocol [2]. Extracted DNA was stored at -80 ºC until qPCR and dehydration. Standards for qPCR were made from a plasmid comprised of *polA* cloned into the pCR 2.1 vector (Invitrogen, Carlsbad, California, USA). One-hundred microliters of DNA was freeze-dried in individual tubes using an ALPHA 1-2 LD Plus freeze dryer (Martin Christ, Germany) and shipped to UNC overnight at ambient temperature.

*RNA oligonucleotide probe design*

In brief, we generated a core set of unique probes that covered all previously called positions of the SS14 and Nichols genomes. We then added probes as needed to cover regions in all other published genomes with <95% sequence homology to the SS14 (GenBank ID: CP004011.1) or Nichols (GenBank ID: CP004010.2) reference strains. Details of the genomes used during the design process are found in **Supplementary Table 2**. Additionally, we increased the tiling density of probes across specific regions of interest, including phylogenetically informative loci and those that encode known or putative outer membrane proteins, to 5x per position (**Supplementary Table 3**). Finally, we excluded any probes predicted to bind human or rabbit genomic DNA.

*Hybridization and library preparation*

After reconstitution, a subset of samples was subjected to ppWGA as described in the manuscript. Prior to hybridization and library construction, genomic DNA (gDNA) or WGA product was diluted in low-EDTA TE buffer (10mM Tris-HCl with 0.1 mM EDTA, pH 8.0) to achieve 20 ng to 200 ng of total DNA in a final volume of 50 µL (**Supplementary Table 4**). Samples were acoustically sheared to 150-200 bp fragments using a Covaris E220 instrument (Covaris, Woburn, MA, USA) and then enriched and prepared for sequencing according to the SureSelect$^{XT\ HS}$ Target

Enrichment System for Illumina Paired-End Multiplexed Sequencing Library protocol. Sequencing was performed at the UNC High-Throughput Sequencing Facility using the MiSeq platform with 150 bp paired-end reads.

*Sequence read alignment and variant calling*

Raw reads were downloaded from the Sequence Read Archive (SRA) database, except for a subset of strains for which raw reads were unavailable (**Supplementary Table 13**). For this subset, we simulated FASTQ files from the publicly available FASTAs using InSilicoSeq's MiSeq model, harvesting 9 million reads per sample. We also simulated FASTQ files from 18 "complete" genome FASTA files. We simulated data from these FASTA files instead of using publicly available raw FASTQ files because these genomes had been previously finished using Sanger sequencing to correct sequencing errors and/or fill gaps. These 18 simulated FASTQs were used as temporary inputs for *GATK*'s HaplotypeCaller to increase the accuracy of SNV calls but were not included in the final whole-genome phylogenetic analysis.

Sequencing reads were pre-processed with Trimmomatic and BBmap to remove human reads, then aligned to the Nichols strain (CP004010.2) or the SS14 strain (CP004011.1) using *BWA MEM* separately [3]. *PICARD* tools and *GATK* were used to perform PCR deduplication and indel realignment, respectively [4, 5]. Low-quality mapping (MAPQ<40), possible cross-mapping artifacts, and genomic regions biased by mapping of reads generated by NGS were removed from the analyses by using the parameter set previously described by Grillova *et al.* [6]. Variant calling was performed using the *GATK* (version 3.5) HaplotypeCaller utility. We then joint genotyped across samples using *GATK*'s GenotypeGVCFs module. Hard-filters for SNVs were employed to capture high-quality variants as follows: QD < 2.0, MQ < 40.0, FS > 60.0, SOR > 3.0, DP <3, MQRankSum < -12.5, ReadPosRankSum < -8.0. SNV calls were made for SS14-like and Nichols-like strains separately.

*Phylogenetic analysis*

Variant calling data were used to generate consensus sequences for phylogenetic analysis. We restricted analysis to samples with at least 3-fold coverage across >80% of the genome. Eighteen complete and 56 consensus "draft" genome sequences were then processed with whole-genome alignment by *MAFFT[7]*. *Gubbins* was applied to these 74 aligned genomes to identify recombination using an algorithm that iteratively identifies loci containing elevated densities of base substitutions [8]. Indels were removed using *VCFtools* before running the *Gubbins* utility [9]. Recombination regions were replaced with N's among all 74 genomes prior to export to *RaxmL* for tree construction. We created a phylogenetic tree based on 1,000 bootstrapped replicates and visualized the tree using the *ggtree* package in *R* (R Core Team, Vienna, Austria)

[10, 11]. Annotation and classification of the functional effects of variants was performed using *SnpEff* [12].

*Sanger sequencing of select loci for independent strain typing by MLST and assessment of macrolide resistance*

First, we compared our phylogenomic strain typing results to those from a multilocus strain typing (MLST) system recently proposed by Grillova *et al.* [13]. MLST typically involves Sanger sequencing of a small number of phylogenetically informative loci and is commonly used for strain typing of a variety of pathogens. To enable comparison of strain-typing calls based on phylogenomic analysis *vs.* MLST, we amplified and Sanger sequenced *tp0136*, *tp0548* and *tp0705* using nested PCR with primers and reaction conditions as previously described [13]. Amplicons were sequenced bidirectionally, and consensus sequences called using Chromas software (Technelysium Pty Ltd, Australia). Sequences were then analyzed using the PubMLST database (https://pubmlst.org/). In addition, we amplified and Sanger sequenced each of the two TPA 23s rRNA operons using operon-specific primer sets as described previously.

### SUPPLEMENTARY RESULTS

*Mutations in genes associated with penicillin-binding*

We observed several mutations in penicillin-associated genes (**Supplementary Table 9**). Penicillin-binding protein mutations in the *mrcA* gene were observed in all of our Chinese isolates, similar to prior observations[14]. Only the Chinese Nichols-like genome harbored an *mrcA* A506T mutation, which has been observed in recent Nichols-like genomes derived from other locations. The significance of these mutations is unclear given that treatment failure after penicillin has not been reported.

*Confirmation by MLST*

We successfully amplified all three loci in all samples and identified three allelic profiles (1.1.1, 1.1.8 and 3.2.3) among our isolates. Allelic profiles of all TPA genomes from China are shown in **Supplementary Table 11**, including 9 Chinese isolates described previously. Allelic profile 1.1.8 predominated among Chinese isolates and was observed in nine of 15 total Chinese isolates. The MLST results were consistent with the global phylogenetic diversity described in **Figure 2B**, including identification of a unique cluster of three SS14-like Chinese strains (allelic profile 1.1.1) and a Nichols-like strain (allelic profile 3.2.3).

*Macrolide resistance of Chinese isolates*

We successfully amplified both operons of the 23S rRNA genes in all six isolates. Sanger sequencing results confirmed A2058G mutations in both operons of all isolates, with the exception of SMUTp_08, which had A2059G mutations. All of our isolates had genetic evidence of macrolide resistance (**Supplementary Table 12**).

A)

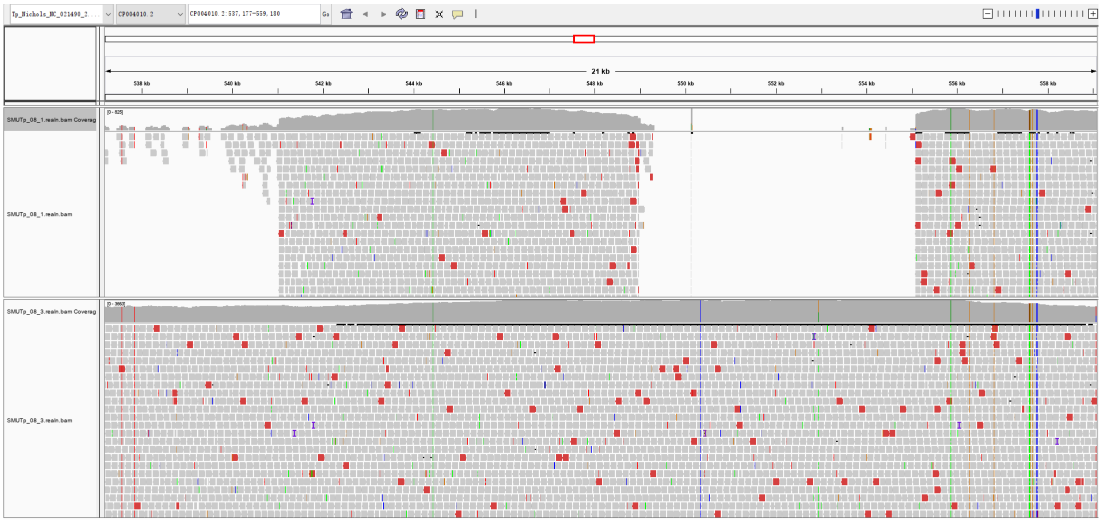

B)

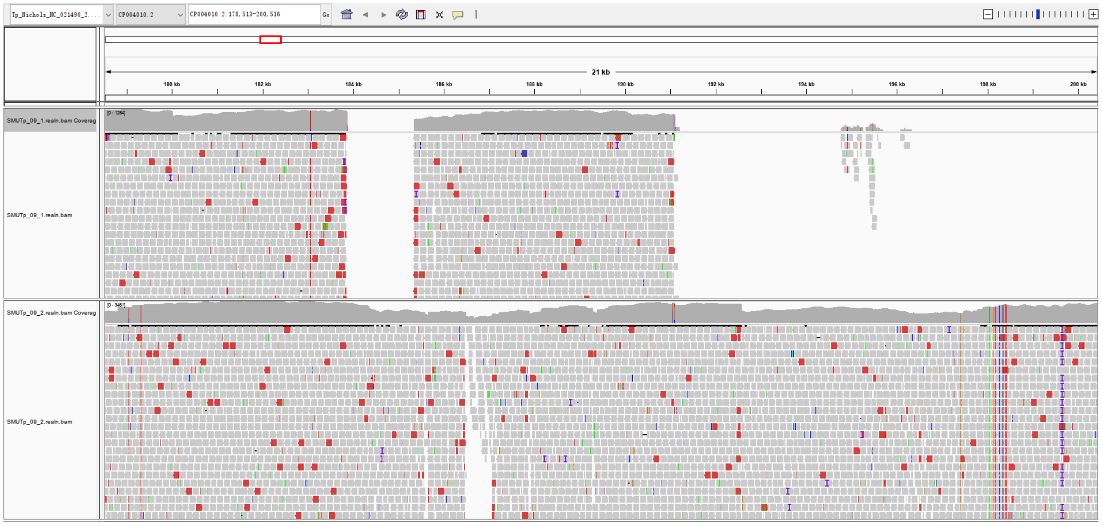

**Supplementary Figure 1.** Sequence read alignments demonstrated islands of coverage observed when a single WGA reaction (top row) was performed prior to hybrid selection, compared to more even coverage observed when five WGA reactions (bottom row) were performed during ppWGA (bottom row) before hybrid selection. Representative screenshots of read alignments for SMUTp_08 (A) and SMUTp_09 (B) from Integrative Genomic Viewer (IGV, Broad Institute and the Regents of the University of California) are shown.
